# The translation inhibitor cycloheximide affects ribosome profiling data in a species-specific manner

**DOI:** 10.1101/746255

**Authors:** Puneet Sharma, Benedikt S. Nilges, Jie Wu, Sebastian A. Leidel

## Abstract

Ribosome profiling provides genome-wide snapshots of translation dynamics by determining ribosomal positions at sub-codon resolution. To maintain this positional information, the translation inhibitor cycloheximide (CHX) has been widely used to arrest translating ribosomes prior to library preparation. Several studies have reported CHX-induced biases in yeast data casting uncertainty about its continued use and questioning the accuracy of many ribosome profiling studies. However, the presence of these biases has not been investigated comprehensively in organisms other than *Saccharomyces cerevisiae*. Here, we use a highly standardized and optimized protocol to compare different CHX-treatment conditions in yeast and human cells. Our data suggest that unlike in *S. cerevisiae*, translating ribosomes in human cells are not susceptible to conformational restrictions by CHX. Furthermore, CHX-induced codon-specific effects on ribosome occupancy are not detectable in human cells nor in other model organisms including *Schizosaccharomyces pombe* and *Candida albicans*. In fact, we find that even in *S. cerevisiae* most biases can be avoided by omitting CHX pre-treatment, indicating that other parameters of library generation contribute to differences between ribosome profiling experiments. Together our findings provide a framework for researchers who plan their own ribosome profiling experiments or who analyze published datasets to draw judicious conclusions.

## Introduction

Protein synthesis is a multilayered process involving a plethora of factors that need to be coordinated in response to cellular and environmental cues (1). The ribosome, an intricate cellular machine, is central to this process as it translates the genetic information encoded in the messenger RNA (mRNA) into a peptide sequence (2, 3). Even though work from many labs has provided a detailed understanding of the biochemical and kinetic aspects of translation — often using specific model substrates — technical limitations have prevented us from understanding its regulation at a global level.

However, this has changed with the advent of ribosome profiling, a breakthrough method that enables the quantitative analysis of transcriptome-wide translation by determining ribosome positioning at sub-codon resolution (4). It has allowed us to comprehend core properties of translation (5), factors governing translational dynamics (6–8) and to characterize individual translation events (9–13). Ribosome profiling is based on deep sequencing of mRNA footprints that are protected from nucleolytic degradation by ribosomes (4, 14). However, despite its use across many species, cell lines and tissues, ribosome profiling studies have reached diverging conclusions when addressing equivalent topics and biological samples, such as the identification of ORFs in zebrafish (9, 10, 15), the influence of wobble base-pairing on elongation speed (16, 17), or the relationship between elongation rates and tRNA abundance (13, 18).

This is in part caused by modifications to the original protocols to match the experimental needs of individual labs (4, 10, 19–22). These differences include the methods of sample harvesting (19, 23), nuclease digestion to generate ribosome footprints (24), preparation of libraries by circularization or dual ligation of linkers and more (4, 9, 22, 25–27). The presence of these small but crucial differences may change the representation of *in vivo* translational dynamics making direct comparisons between datasets challenging. When harvesting yeast, cultured mammalian cells or tissue samples, two key elements likely influence the outcomes of ribosome profiling experiments: The speed of harvesting and the use of translation inhibitors. To faithfully capture the *in vivo* conformation and positioning of translating ribosomes, the timespan between the onset of harvesting and flash-freezing of samples should be kept to a minimum (23). Another failsafe against undesired elongation *in vivo* and in the lysate, is the pre-treatment with the translation inhibitor cycloheximide (CHX) used since the original protocols in yeast (4) and mammalian cells (28). CHX is a small molecule that inhibits translation elongation by binding to the ribosomal E-site (29, 30). Different studies, in particular in *Saccharomyces cerevisiae*, have reported biases triggered by CHX pre-treatment (31–36). For example, CHX appears to reversibly interact with ribosomes, allowing them to move away from their initial position on mRNA (33). This movement is influenced by codon identity, therefore, altering the outcomes of studies that require codon-level resolution for occupancy and translation speed measurements (33). Furthermore, high concentrations of CHX inhibit elongation, but not initiation (29) leading to ribosome accumulation at the start codon (5). CHX might even stabilize a particular conformation of ribosomes thereby biasing ribosome profiling data towards RNA or peptide features that facilitate such a conformation (32).

Despite those concerns, the addition or omission of CHX has not been systematically compared in experiments using standardized conditions to the best of our knowledge. Most analyses compared published ribosome profiling data generated by different labs or varied several conditions simultaneously (33). Therefore, some past conclusions on the effects of translation inhibitors are likely affected by parameters other than CHX usage. Furthermore, the effect of CHX on ribosome profiling has been predominantly studied in *S. cerevisiae* and we know little about its impact in vertebrates and other model organisms. We, therefore, conducted a comprehensive analysis of the effect of CHX on ribosome profiling experiments in different yeasts and mammalian cells. Libraries were generated with a highly standardized and optimized protocol to maximize comparability and to deduce CHX-specific biases.

We describe a protocol that confidentially maps high numbers of the ribosome footprints to the ribosomal A-site without the need to subtract ribosomal RNA (rRNA). Implementing this protocol, we systematically assessed the impact of CHX on ribosome profiling experiments and found that human ORFs show a slower elongation rate in the first 15~150 codons irrespective of CHX treatment. Furthermore, the drug treatment does not affect translation at the gene level. Importantly, in contrast to yeast, ribosomes of humans and other common model organisms are insensitive to the CHX-induced alteration of apparent codon-specific translational velocity. These findings show that CHX affects the measurements of translation dynamics in a species-specific manner. Therefore, concerns against the use of cycloheximide in ribosome profiling are not warranted in most experimental systems and even in baker’s yeast the careful use of CHX does not perturb most types of analyses. However, the observed differences in overall parameters of ribosome profiling libraries emphasize the need for better standardization beyond the use of CHX.

## Results

### Stronger digestion of the cell lysate improves footprint size distribution

Translating ribosomes protect mRNA from nuclease digestion during ribosome profiling experiments, giving rise to a distinct footprint size distribution (14). If nuclease digestion is stringent, unprotected nucleotides are efficiently trimmed leading to a characteristic three-nucleotide periodicity of mapped reads (4). The larger the fraction of footprints that map to the correct reading-frame of the ORFs the higher the confidence, with which ribosomal A-, P- and E-sites can be assigned to specific positions within the footprints. Therefore, we initiated our analysis by establishing a robust ribosome profiling protocol that minimizes biases and yields a majority of footprints in the correct reading frame. We harvested and lysed yeast and HEK 293T cells by rapid cryogenic freezing and studied the effect of different digestion strengths on library quality by varying RNase I concentration. In yeast, stronger RNase I digestion increased the fraction of 28 nt footprints (69% of all reads at 600 U digestion) and improved in-frame reads (frame 0) of 28 nt reads to 90% (Supplementary Figure 1A and 1B). A similar effect was observed in human cells with footprint-distribution centered around 30 nt (Supplementary Figure 1C). Interestingly, stronger digestion of human samples provided the additional benefit of decreased rRNA contamination in the libraries (Supplementary Figure 1D). Based on this analysis, we generated ribosome profiling libraries from lysates that were treated with high amounts of RNase I (600 U for yeast and 900 U for HEK 293T cells lysates).

### Cycloheximide differentially affects human and yeast ribosomes

CHX induced biases have been reported in yeast ribosome profiling (31, 33–35). However, even in yeast most analyses compared datasets that were not generated in parallel, and no study has systematically investigated the effects of CHX at the different steps of sample preparation in human cells. Hence, we directly compared three treatments: First, we briefly incubated HEK 293T cells in medium containing CHX prior to lysis and included the drug in the lysis buffer (+/+). Second, the cells were subjected to CHX only in the lysis buffer (−/+). Finally, we lysed the cells without pre-incubation and in a buffer devoid of the drug (−/−). All experiments were performed in parallel and cells were subsequently lysed cryogenically in liquid nitrogen to limit ribosome run-off. For inter-species comparisons, we harvested yeast via rapid filtration and flash froze the cells using the same three types of CHX treatment (Figure 1A).

**Figure 1:**
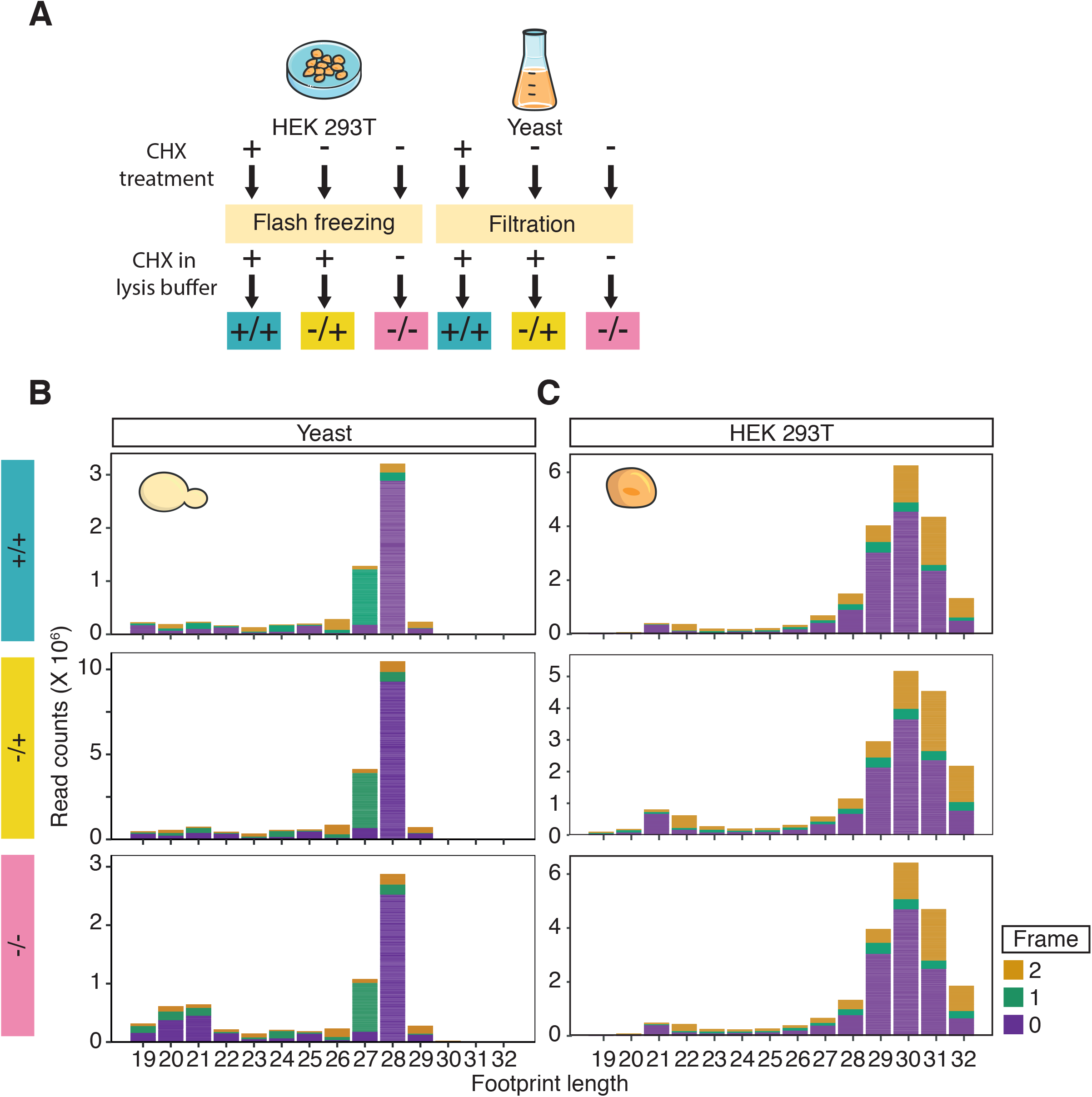
Cycloheximide (CHX) differentially affects footprint length in yeast libraries. A) Schematic overview of harvesting and CHX-treatment conditions used in this study. B) Histograms showing the influence of CHX on footprint length and the reading frame in yeast libraries. C) Same as B) for HEK 293T cells. Values stem from three biological replicates. The reading frame is indicated by color: 0 (purple), 1 (green) and 2 (yellow).

CHX is thought to stabilize ribosomes in a specific rotated conformation during the elongation stage, yielding 28-30 nt footprints (5, 32). In the absence of CHX, 20-22 nt footprints can be detected in libraries, possibly representing a non-rotated ribosomal conformation (32). To detect footprints representative of both conformations, we isolated RNase-protected fragments between 18 and 32 nt. In agreement with previous reports (32), we observed that the omission of CHX in yeast leads to a mixture of two distinct footprint sizes (20-21 nt and 27-28 nt; Figure 1B). The short footprints were specific to (−/−) libraries, pointing to a CHX-mediated effect, that prevents the recovery of short footprints, even if it is only added after lysis. Surprisingly, we observed that human ribosomes contain short footprints (21-22 nt) irrespective of CHX treatment (Figure 1C). To exclude that the CHX treatment was too weak to stabilize ribosomes in a conformation yielding only long footprints, we increased incubation time and CHX concentration. However, the short footprints persisted regardless of the duration and strength of CHX treatment (Supplementary Figure 1E), suggesting that CHX affects human and yeast ribosome profiling experiments differently. In agreement with our observation, short footprints in human cells can also be observed in a recently published dataset that uses CHX treatment for up to 24 hours (Supplementary Figure 1F) (37). This shows that the use of CHX in ribosome profiling has to be assessed for each organism and prompted us to analyze different parameters of translation, focusing on larger footprints for better comparability with previous publications.

### Translation ramp and ribosomal occupancy are independent of CHX

Ribosome profiling data can be used to infer elongation rates along a transcript because local footprint densities reflect the residence times of ribosomes. Several studies in yeast have observed an increase in ribosome density in the first ~200 codons of ORFs. This phenomenon is called the ‘5’ translation ramp’ and is thought to represent slow elongation speed in this region (38). In contrast, a study in murine embryonic stem cells concluded that a similar ramping effect is not observed in vertebrate cells (5). To elucidate the effect of CHX on the onset and termination of translation, we analyzed footprint density around annotated start and stop codons. Consistent with previous reports, we observed increased ribosome density in the first ~15 codons of well-expressed genes in yeast and also in human (+/+) libraries and to a lesser extent in the (−/+) and (−/−) libraries (Figure 2A). Importantly, we found an enrichment of ribosomes from 15 to ~150 codons irrespective of the drug treatment in both species. The scale of this effect and the position of the maximum footprint density (~60 codons in humans, and ~40 codons in yeast) are independent of gene-translation levels and gene length (Supplementary Figure 2; and data not shown). This shows that even though CHX leads to an accumulation of footprints immediately downstream of the start codon, the translation ramp itself is not induced by CHX, but a genuine feature of translation in yeast and humans. Finally, we observed an accumulation of ribosomal footprints at stop codons in all three conditions, but most prominently in the (−/+) libraries (Figure 2A and 2B). This is consistent with reports in murine and yeast cells (5, 39, 40).

**Figure 2:**
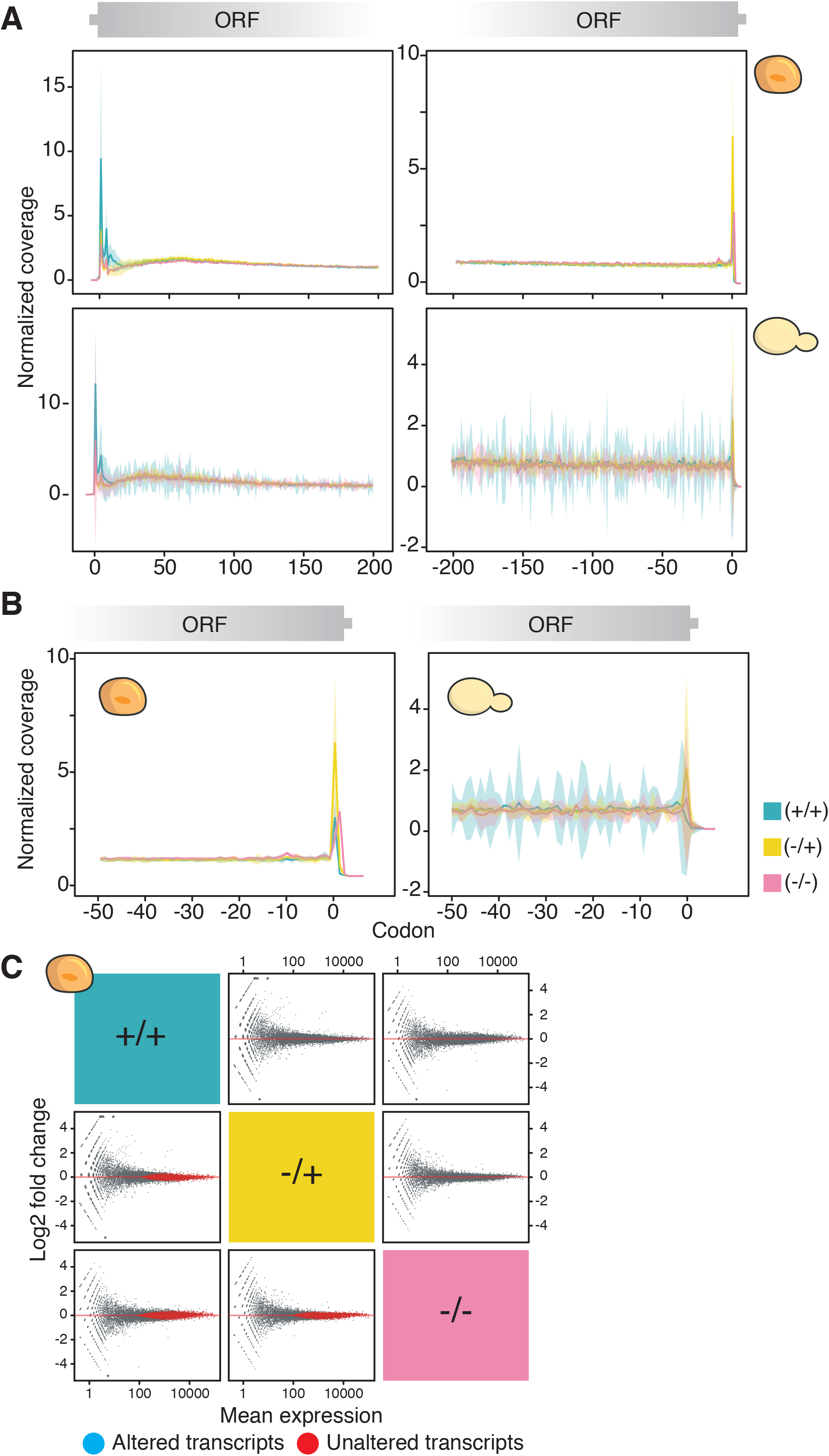
Metagene analysis shows CHX-independent ribosome accumulation 15-150 codons downstream of the start codon and mammalian ribosome occupancy at the ORF level is impervious to CHX treatment. A) Normalized ribosomal A-site coverage for the first 200 (left) and last 200 (right) codons in HEK 293T cells (top) and yeast (bottom) in highly expressed genes (>64 reads). Shaded areas represent confidence intervals (n = 3). B) Like in A) but showing the normalized ribosomal A-site coverage of the last 50 codons in HEK 293T cells (left) and yeast (right). CHX-treatment conditions are indicated by color (+/+) (blue), (−/+) (yellow), (−/−) (pink). C) Differential ribosome occupancy across ORFs in HEK 293T cells across inhibitor treatments determined with DESeq2 (41). ORFs were tested for differential translation (top right, adjusted p-value ≤ 0.05, log2 fold change threshold = 0.5) and for unaltered translation (bottom left, adjusted p-value ≤ 0.05). Significant ORFs are highlighted in red.

Our analysis of the translation ramp implied, that CHX does not induce a global effect on ribosome coverage. However, transcripts enriched for specific codons might be susceptible to CHX-induced alteration of apparent translation velocity, prompting us to analyze whether individual transcripts display altered ribosome occupancy when using CHX. CHX treatment of yeast and murine cells was previously shown to not affect expression measurements at the transcript level (5, 33, 40). Since we found that the effects of CHX appear to be species specific, we analyzed its impact in human cells. We counted uniquely mapped ribosome footprints across the human transcriptome and determined differential abundance between (+/+), (−/+) and (−/−) libraries using DESeq2 (41) (Figure 2C). We did not observe transcripts that were significantly altered by CHX treatment (Figure 2C; top right). To exclude that this was caused by technical variability of the sequencing libraries we tested for transcripts that were not altered with statistical certainty (Figure 2C; bottom left). The analysis yielded 7109 (+/+ vs. −/+), 8056 (−/+ vs. −/−) and 7595(+/+ vs. −/−) transcripts that were not affected out of approximately 14000 detected ones. These high numbers exemplify the statistical power of this approach, that would have detected CHX-dependent alterations, if they existed. Thus, human cells do not display altered ribosome occupancy at the transcript level upon CHX treatment and translation levels can be compared irrespective of CHX use.

### Cycloheximide does not affect codon level ribosome occupancy in human cells

CHX arrests translation by binding to the ribosomal E-site thereby locking the ribosome in the rotated state (32). However, binding kinetics might depend on the decoded codons leading to a mis-representation of specific codons as described for CGA and CGG in yeast (33). This would be perceived as an alteration in decoding kinetics of specific subsets of codons, but may remain undetected when quantifying occupancy across transcripts consisting of hundreds of codons. To test for the CHX-dependent enrichment of codons, we calculated transcriptome-wide A-site codon occupancy in our human and yeast data (Figure 3). Correlations between HEK-libraries were excellent (R^2^ ≥ 0.94 between all conditions (Figure 3; top right)), showing that A-site occupancy remains unaltered by the use of CHX in ribosome profiling experiments. In contrast, yeast samples with and without pre-treatment correlated much weaker (R^2^ (+/+ vs. −/+) = 0.48 and R^2^ (+/+ vs. −/−) = 0.54 (Figure 3; bottom left)) and the addition of the inhibitor in the lysis buffer had a lower impact on A-site occupancy (R^2^ (−/− vs. −/+) = 0.86). Importantly, this effect is not specific to our dataset, but is also apparent in other studies that varied the moment of CHX addition (8, 31, 32, 42–44) (Supplementary Figure 3). Hence, the use of CHX in ribosome profiling needs to be viewed in a species-specific manner and conclusions made in yeast cannot necessarily be transferred to humans. Furthermore, even in yeast using CHX in the lysis buffer without pretreatment of the culture leads to only minor differences.

**Figure 3:**
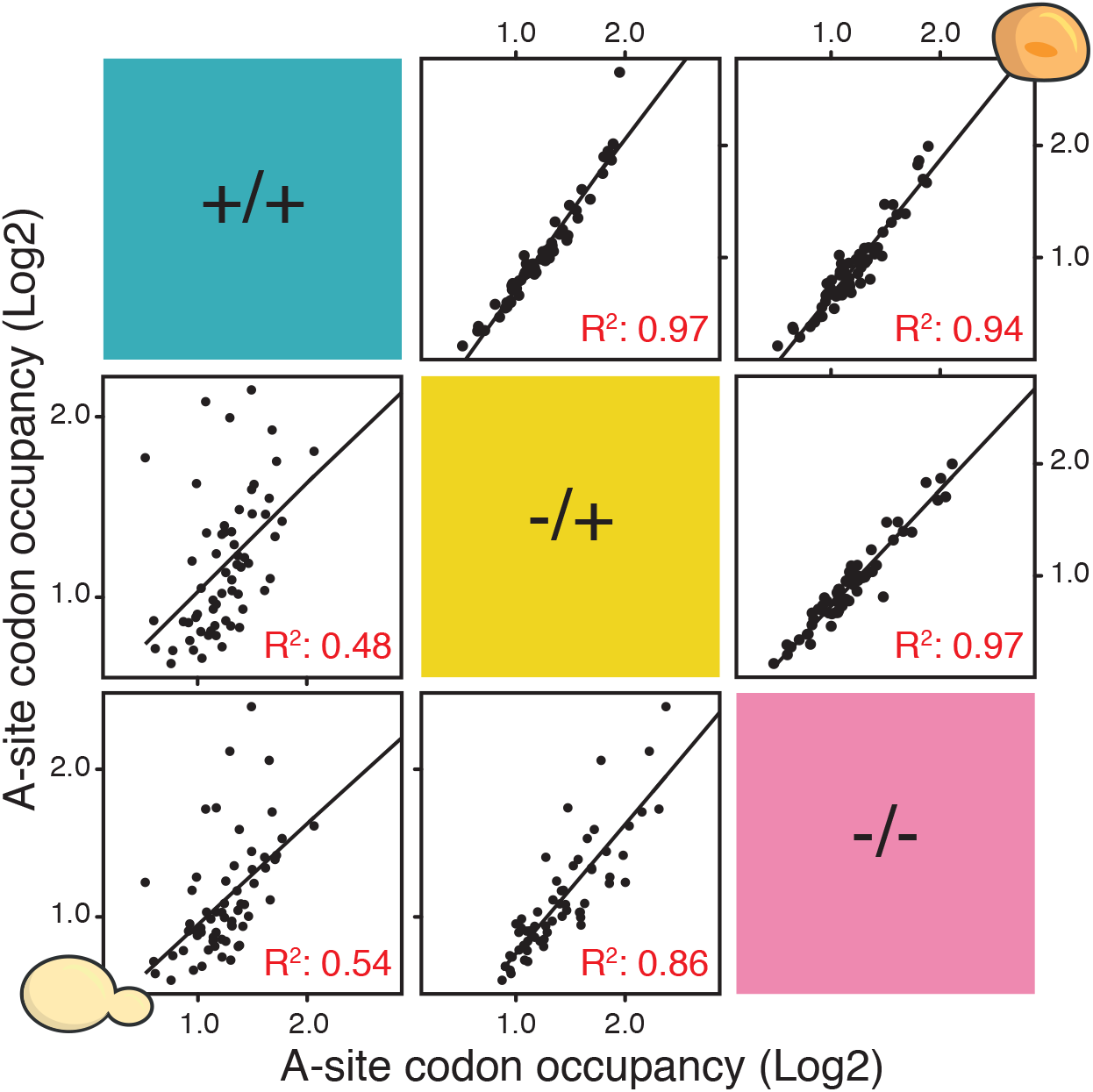
Cycloheximide (CHX) does not alter mammalian ribosome occupancy at the codon levels. Comparison of transcriptome-wide A-site codon occupancy in HEK 293T cells (top right) and yeast (bottom left) across inhibitor treatments (mean, n=3). Each black dot represents a codon.

### The absence of CHX dependent codon waves is consistent across most species

A consistent pattern of increased ribosome occupancy downstream of specific codons was described in CHX-treated yeast samples (33). This phenomenon — often referred to as ‘waves’ — occur at the rare codons CGA and CGG that are decoded by the essential 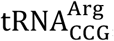, which is expressed from a single gene (45, 46). CGA requires a wobble interaction to be decoded and exhibits a more pronounced wave than CGG, which is the cognate codon of 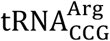. Consistent with this, we observed a wave for CGA and CGG in yeast (Figure 4A and data not shown). We tested whether waves downstream of CGA are a common feature of eukaryotic ribosomes, but did not observe a similar ribosome enrichment in HEK cells (Figure 4B). Since there is no codon in the human transcriptome that is as rare as yeast CGA and decoded by a single tRNA we analyzed wave formation for all codons in the human transcriptome (Supplementary Table 1). Interestingly, we did not observe wave formation for any codon in humans (data not shown). Even for the human UUA codon, which is relatively rare with a low tRNA copy number, we did not detect CHX-induced downstream enrichment of ribosomes. Waves are not only absent from human cells following our protocol, but from all published human ribosome profiling datasets that we analyzed (Figure 4C and Supplementary Figure 4). This effect is not specific to human cells, because waves of ribosome occupancy were generally absent from vertebrate ribosome profiling data (Figure 4D). Neither zebrafish (10) nor mouse (5) featured a wave downstream of any codon. Finally, to investigate whether the altered translation at rare codons is a feature of yeasts we generated ribosome profiling libraries from CHX treated *Schizosaccharomyces pombe* and *Candida albicans*. Similar to *S. cerevisiae*, CGA and CGG are rare codons in these species and are decoded by only one tRNA each (*S. pombe*) or rely on a single tRNA for both codons (*C. albicans*). Surprisingly, the data from these two yeasts was free of waves at CGA and CGG or other codons (Figure 4E). This suggests that the slowdown of translation following these codons is highly specific to baker’s yeast and that the use of CHX is, therefore, only a concern when studying translation in *S. cerevisiae*, but not in other widely used eukaryotic model organisms.

**Figure 4:**
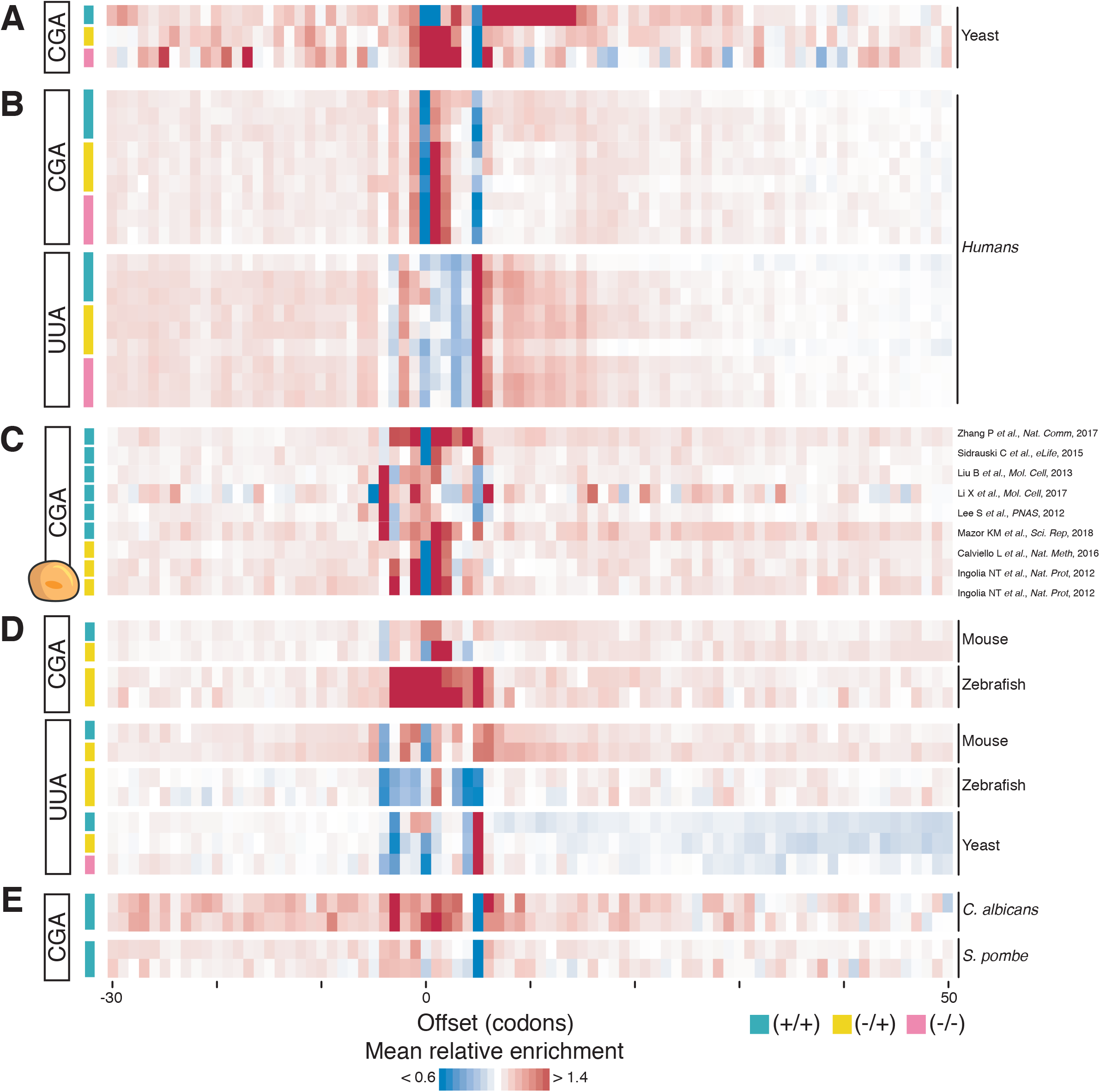
Cycloheximide (CHX) pre-treatment of vertebrate samples does not alter ribosome occupancy downstream of rare codons. A) Transcriptome-wide ribosome enrichment profiles of yeast cells surrounding CGA codon with different CHX treatment methods according to Hussmann and colleagues (33). B) Same as A) for HEK 293T cells surrounding CGA and UUA codons with different CHX treatment. C) Same as A) for the CGA codon on published human ribosome profiling datasets (19, 20, 48–53). D) Same as A) for the CGA and UAA codons from E14 mouse embryonic stem cells (5), zebrafish embryos (10) and wild-type yeast (this study). E) Same as A) for the CGA codon in *C. albicans* and *S. pombe*. CHX treatment conditions are indicated by color: (+/+) (green), (−/+) (yellow), (−/−) (pink).

### Variability in harvesting and footprinting may contribute to the poor correlation in similar samples

Ribosome profiling data from different laboratories correlated poorly even within the same CHX-treatment regime (Figure 5 and Supplementary Figure 3). A closer inspection revealed a large variability in the cell harvesting, gradient centrifugation, the amount of cell lysate, RNase I digestion temperature, digestion time and the library-preparation methods. These details are often overlooked when comparing ribosome profiling experiments and can have profound effects on ribosome occupancy (Supplementary Figure 5). However, our analysis suggests that they are more critical than they appear at first glance and far outweigh the effects of CHX-treatment.

**Figure 5:**
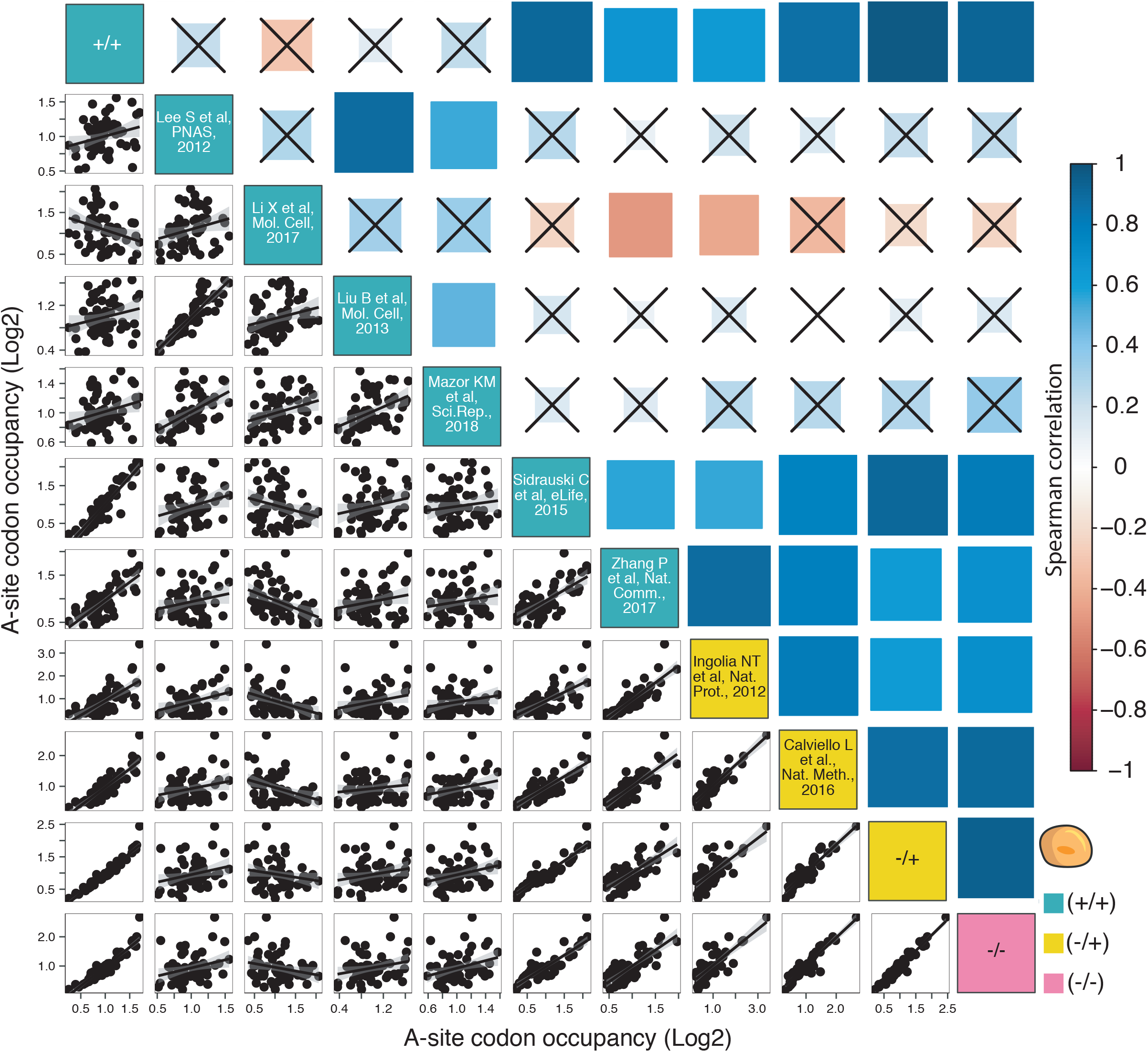
Cycloheximide (CHX) does not explain poor correlation of various datasets in humans. Linear regression analysis of transcriptome-wide A-site codon occupancy across data from this study and published datasets for human cells with different inhibitor treatments (19, 20, 48–53). Each black dot represents a codon. CHX-treatment conditions are indicated by color: (+/+) (green), (−/+) (yellow), (−/−) (pink). Size of the box indicates p-value. Correlations with a p-value > 0.05 are crossed out.

## Discussion

Caution has been advised when using CHX in ribosome profiling experiments based on reports of CHX induced biases in yeast (31–33, 47). This has led some researchers to generally question results of studies that utilize CHX and may have even prompted them to invest time and resources to conduct experiments without the inhibitor to confirm previous findings. To the best of our knowledge, we have conducted the first systematic analysis to determine the effects of CHX at the different stages of library preparation in parallel experiments to assess its impact. Importantly, when analyzing yeast and human libraries, we find the effect of CHX on the data to be species dependent. Human ribosome profiling libraries generated by rapid lysis and without prolonged exposure to CHX prior to lysis are devoid of reported CHX-mediated biases. We did not find significant differences in overall translation levels, codon enrichment and translation dependent features like the translational ramp, a finding that we verified by analyzing published datasets (8, 19, 20, 22, 48–53).

Both human and yeast cells exhibit a translational ramp. Though initially hypothesized as induced by CHX pre-treatment we find it to be independent of CHX treatment emphasizing that the translational ramp is indeed a genuine biological feature (40). Multiple factors were proposed to trigger the translational ramp, such as the presence of rare codons with low cognate tRNA availability, mRNA secondary structure or the interaction of nascent polypeptides with the ribosomal exit tunnel (38, 54). Even though its mechanistic reasons are still debated we were able to exclude that CHX treatment, gene length or transcript expression levels are primary contributing factors (Supplementary Figure 2A). It is an exciting possibility that the formation of the ramp prevents the likelihood of ribosome collisions, which activate ribosomal quality control (RQC) and trigger mRNA decay (55).

The codon-mediated alteration of translation speed in CHX treated baker’s yeast appears to affect rare codons with low levels of cognate tRNA, such as CGA and CGG (33). There is no codon in commonly used vertebrate models with a similarly low codon frequency and tRNA gene copy number like CGA in yeast, suggesting that these organisms are less likely to be affected by CHX treatment. Consistently, we found no evidence of ribosome enrichment downstream of any codon in zebrafish, mouse and human ribosome profiling datasets. We would like to point out that it will require even more replicates and deeper sequencing to detect subtle effects in specific ORFs or whether alteration in translation speed at codon combinations may be induced by CHX. However, even in *S. cerevisiae* wave formation likely reflects a genuine structural feature of its ribosomes. Neither a low codon frequency nor tRNA copy number can explain CHX-mediated waves as shown by the comparison with *S. pombe* and *C. albicans*.

The marked difference between baker’s yeast and other species might arise from two factors: First, differences in the uptake and turnover of CHX and second, the structure of the yeast ribosome and how CHX interacts with it. We excluded the possibility that slower uptake of CHX in human cells might contribute to our findings by exposing cells to higher concentrations CHX or by prolonging the incubation time (Supplementary Figure 1E). Yeast and human ribosomes are structurally highly similar at the catalytic core yet they exhibit differences. For example, human ribosomes contain longer RNA-expansion segments (56) and an additional large subunit protein, L28e (57). These long expansion segments are highly mobile, which could alter the accessibility of CHX and RNase I and explain different footprint properties. A recent study reported that the binding site of the piperidine-2,6-dione moiety of CHX in human ribosomes is similar to yeast, but that the ligand adopts a different conformation and binding mode (58). This may facilitate a more stable binding of CHX to human ribosomes leading to a better preservation of their localization on the mRNA. CHX might even associate with human ribosomes that are in the rotated conformation, which would explain the presence of short ribosome footprints in (+/+) samples. In yeast, short footprints are thought to be the product of ribosomes in the rotated state, while the non-rotated state is stabilized by CHX binding (32). Such subtle species-specific differences may underlie some of the observed differences between mammalian and yeast ribosome profiling data in this study. We conclude that the CHX effect in yeast is most likely triggered by subtle structural differences in mRNA-ribosome interactions at particular codons. These differences affect CHX-ribosome interactions. To disentangle the molecular mechanisms will necessitate a detailed analysis of CHX-binding to ribosomes in different organisms.

While we cannot fully exclude that some of the observed discrepancy between *S. cerevisiae* and other species reflects differences in the translation activity of the cell extracts following lysis, we consider this an unlikely scenario. First, we do not observe codon waves in *S. pombe* or the closely related *C. albicans* using the same lysis protocol. Second, the lysis procedure of human cells takes longer than the fast filtering and flash freezing of yeast. This would allow CHX to interact with human ribosomes for a longer timespan than during yeast lysis.

While certainly detectable, the biases introduced by CHX in yeast libraries are less severe than proposed. They do not affect global gene translation level, the translation ramp and most of the individual codons. Therefore, most datasets can be analyzed by cautiously excluding the codons surrounding the start and the stop codon from an analysis. Naturally, prudence will be required when CGA and CGG codons are concerned. However, this may not be specific to ribosome profiling since these two codons also trigger phenotypes in assays that rely on the use of reporter constructs (59–62). By expressing constructs that contain such rare codons the scarcity of tRNA required for their decoding will be increased. Therefore, neither reporter assays nor ribosome profiling experiments should be performed without the proper controls and verifications by orthogonal methods.

While analyzing ribosome profiling data from various studies, we observed discrepancies in general library properties such as fragment length, the extent of frame information and the level of contamination with rRNA or small RNAs (Supplementary Figure 5). Such differences can result from diverging library preparation protocols at the levels of lysis, digestion and size selection and will likely affect the analysis of translation dynamics. Furthermore, we noticed that crucial information is sometimes lacking in the methods section of publications. Therefore, we urge the community to begin standardizing the ribosome profiling protocol and to set minimum standards for reporting how libraries are prepared. This includes (I) the exact method of lysis, (II) the concentration of nucleases relative to the amount of nucleic acids, (III) duration and temperature of footprinting and (IV) what footprint sizes were selected for library preparation. A mere citation of the initial ribosome profiling studies (4, 19) does not provide enough information. However, knowing the details of individual experiments will allow the community to select, which published datasets and translational features can be directly compared.

### Box 1

#### A note on sample harvesting and reporting sample preparation procedure

Since the seminal publication 10 years ago the experimental protocol for ribosome profiling has steadily evolved without consensus on the ideal procedure or how to report its application. Theoretically, a key to faithfully capturing the *in vivo* conformation and positioning of translating ribosomes is the timespan between the onset of harvesting and freezing of the samples. For unicellular organisms like yeast and *Escherichia coli* that grow as a cell suspension, two methods of harvesting have been reported in the literature. The first method uses rapid cooling of the cell suspension followed by centrifugation and flash freezing (23). Alternatively, cells are subjected to rapid filtration and flash frozen (4, 40). The former method risks eliciting a starvation response, and, therefore, ribosome run-off, in the cells due the centrifugation induced stress (63) and is more time consuming compared to the latter methods. Similarly, adherent mammalian cells are harvested and lysed in various ways. The first application of ribosome profiling in vertebrate cells resuspended the cell pellet in ice-cold lysis buffer followed by centrifugation and flash freezing (28). Most recent protocols, however, avoid cell pelleting and use either lysis with or without freezing for rapid harvesting of cells with intact ribosomes (19).

## Materials and Methods

### Yeast harvesting and footprinting

Overnight cultures of wild-type yeasts in the BY4741 (*S. cerevisiae*), 972 (h^−^) (*S. pombe*) and SN87 (*C. albicans*) backgrounds were diluted and grown to mid-exponential phase (0.4 ~ OD_600_). For (+/+) samples, CHX was added to a total concentration of 100 μg/ml and cultures were gently agitated for 1 min at 30 °C. Cells were rapidly harvested by vacuum filtration through a 0.45 μm cellulose nitrate filter (GE Healthcare) and immediately flash-frozen. Samples were mechanically lysed under cryogenic conditions in a Freezer-Mill (SPEX SamplePrep) with 2 cycles at 5 CPS interspersed by 2 min of cooling. Lysates were thawed in lysis buffer (20 mM Tris-HCl pH 7.4, 5 mM MgCl2, 100 mM NaCl, 1 % Triton, 2 mM DTT) containing 100 μg/ml CHX for (+/+) and (−/+) samples and clarified by two rounds of centrifugation (5 min; 4 °C; 10,000 g). Unless specified otherwise, 10 A260 units of cleared lysates were digested with 600 U Ambion RNase I (ThermoFisher) for 1 h at 22 °C and the reaction was inhibited by the addition of 15 μl SuperaseIn (ThermoFisher). Monosomes were isolated from the sucrose gradients according to (8). Finally, libraries were generated as described before (19), using 3’-adapters that were randomized at the first 4 positions of the 5’ end to minimize potential ligation biases (22).

### HEK 293T cell harvesting and footprinting

For the (+/+) condition, cultured HEK 293T cells were incubated with medium containing 100 μg/ml CHX for 1 min, washed with ice cold PBS and flash frozen in liquid nitrogen. The dish was swiftly transferred to ice and 400 μl lysis buffer (10 mM Tris-HCl pH 7.5, 100 mM NaCl, 5 mM MgCl2, 1 *%* Triton X-100, 0.5 mM DTT, 0.5 % deoxycholate (w/v) and 100 μg/ml CHX) was dripped onto the cells. The cells were harvested on ice by scraping once the lysis buffer was thawed. Cells for (−/+) and (−/−) conditions were harvested like (+/+), however, CHX pre-incubation was omitted and it was not added to the lysis buffer for (−/−). Samples were clarified by centrifugation (5 min; 4 °C; 10,000 g). Unless specified otherwise, 10 A260 units of cleared lysates were digested with 900 U RNase I (ThermoFisher) for 1 h at 22 °C and the reaction was inhibited by the addition of 15 μl SuperaseIn (ThermoFisher). Monosomes were isolated from the sucrose gradients and libraries were prepared analogous to yeast samples.

### Sequencing, processing and mapping of reads

Ribosome profiling and RNA-Seq libraries were sequenced on Illumina HiScanSQ and NextSeq sequencers. Ribosome profiling reads were processed by clipping the adapter sequence and trimming the 4 randomized nucleotides of the linker using the FASTX-Toolkit (http://hannonlab.cshl.edu/fastx_toolkit, June 2017), version 0.0.13. Processed yeast reads were mapped to tRNA, rRNA, snRNA and snoRNA genes from SGD to remove possible contaminants using bowtie (64) version 1.0.0. Processed reads from human samples were mapped to rRNA using bowtie version 1.2.1.1. Residual reads were uniquely mapped to non-dubious ORFs (sgdGene) using bowtie. Reference ORFs (hg38 UCSC canonical transcripts; mm10 UCSC genes; danRer10; C_albicans_SC5314_version_A22; Pombe_ASM294v2) were extended by 18 nt into the UTRs to allow alignment of footprints from initiating and terminating ribosomes.

The majority of mapped footprints had a length of 29-31 nt, 30-32 nt and 27-29 nt in human, mouse and yeast, respectively. We assigned these mapped footprints to A-site codons according to the frame of the 5’ end of footprints as previously described (4). For footprints with the 5’end in the −1 frame the A-site was defined as position 17-19 and the ones with 5’ ends in the 0 frame as position 16-18.

For ribosome occupancy at the beginning of ORFs, we normalized the coverage of A-site footprints by the average ribosome occupancy of all codons in that gene:

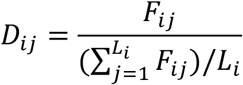

Where *F_ij_* and *D_ij_* are the ribosome footprints and density of positon *j* of gene *i*, respectively. *L_i_* is the length of genes in codons. The average of ribosome densities at the postion *j* is calculated by normalizing to the number of all well-expressed genes (> 64 reads) with an ORF length of at least 200 codons:

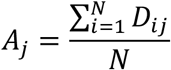

Where *A_j_* is the average of ribosome densities and *N* is the number of genes.

A-site codon occupancy plots (8) and wave plots (33) were generated as described.

## Acknowledgements

We thank Claudia Gräf and Fiona Alings for technical support and sequencing library generation, Namit Ranjan and all members of the Leidel lab for comments on the manuscript and critical discussions. *Schizosaccharomyces pombe* strain 972 (*h*^−^) and *Candida albicans* strain SN87 were kindly provided by Elena Hidalgo and Suzanne Nobel, respectively. This work was supported by the Max Planck Society and DFG [LE 3260/3-1] to S.A.L; P.S., B.S.N. and J.W. gratefully acknowledge fellowships from the Graduate School of the Cells-in-Motion Cluster of Excellence [EXC 1003-CiM].

