## Supplementary Figures for "The translation inhibitor cycloheximide affects ribosome profiling data in a species-specific manner"

Supplementary Information

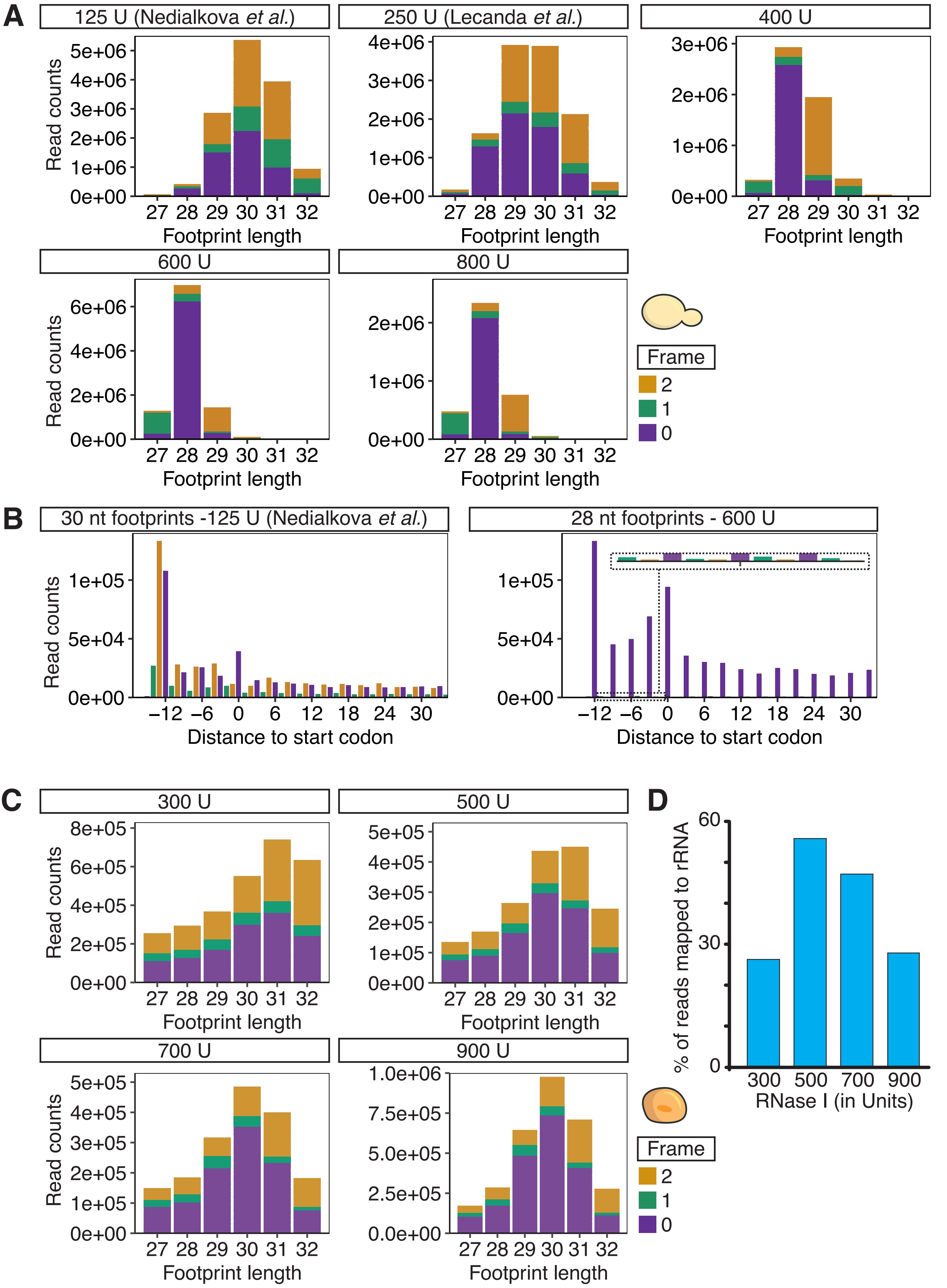


Supplementary Figure 1

E

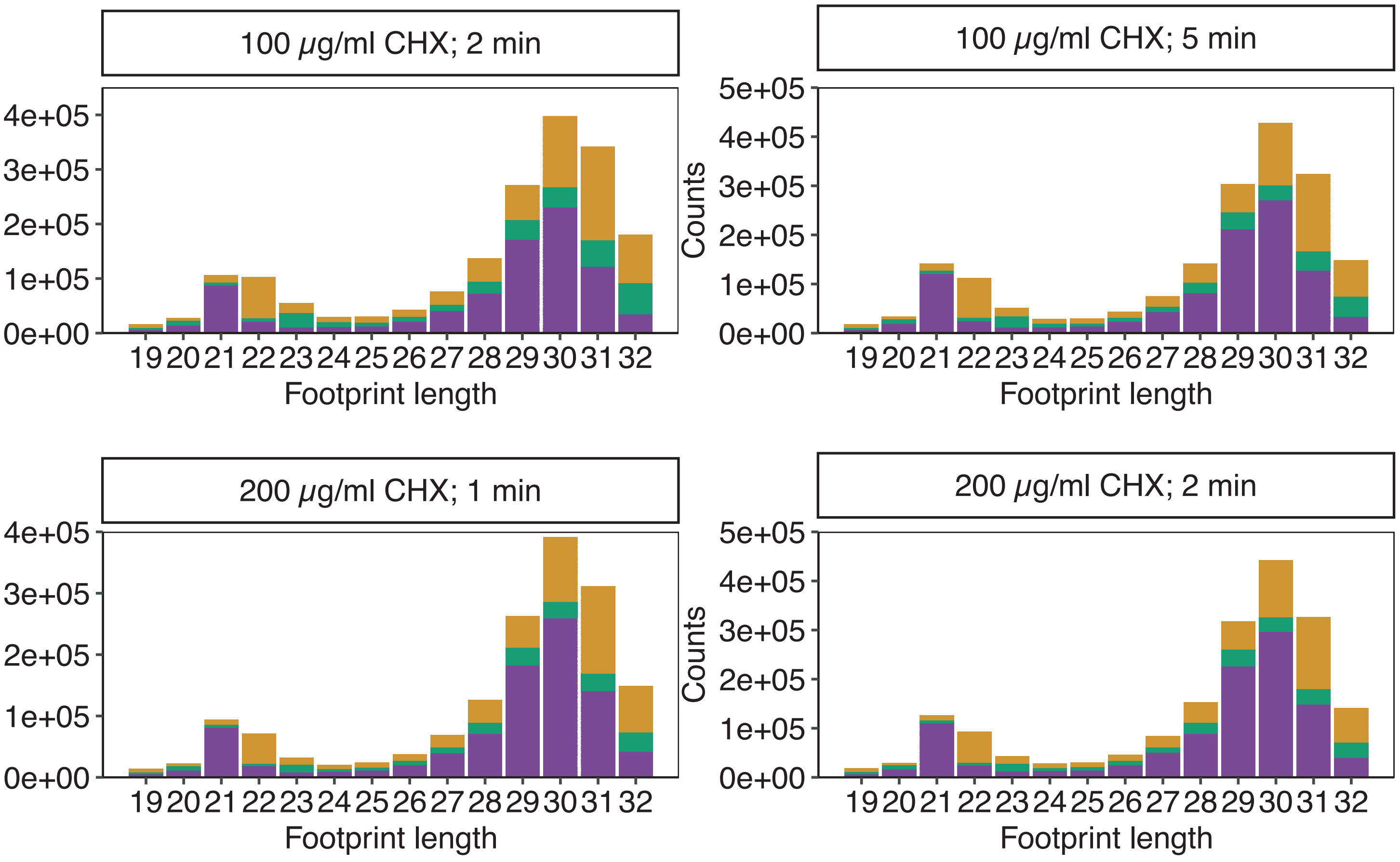


F

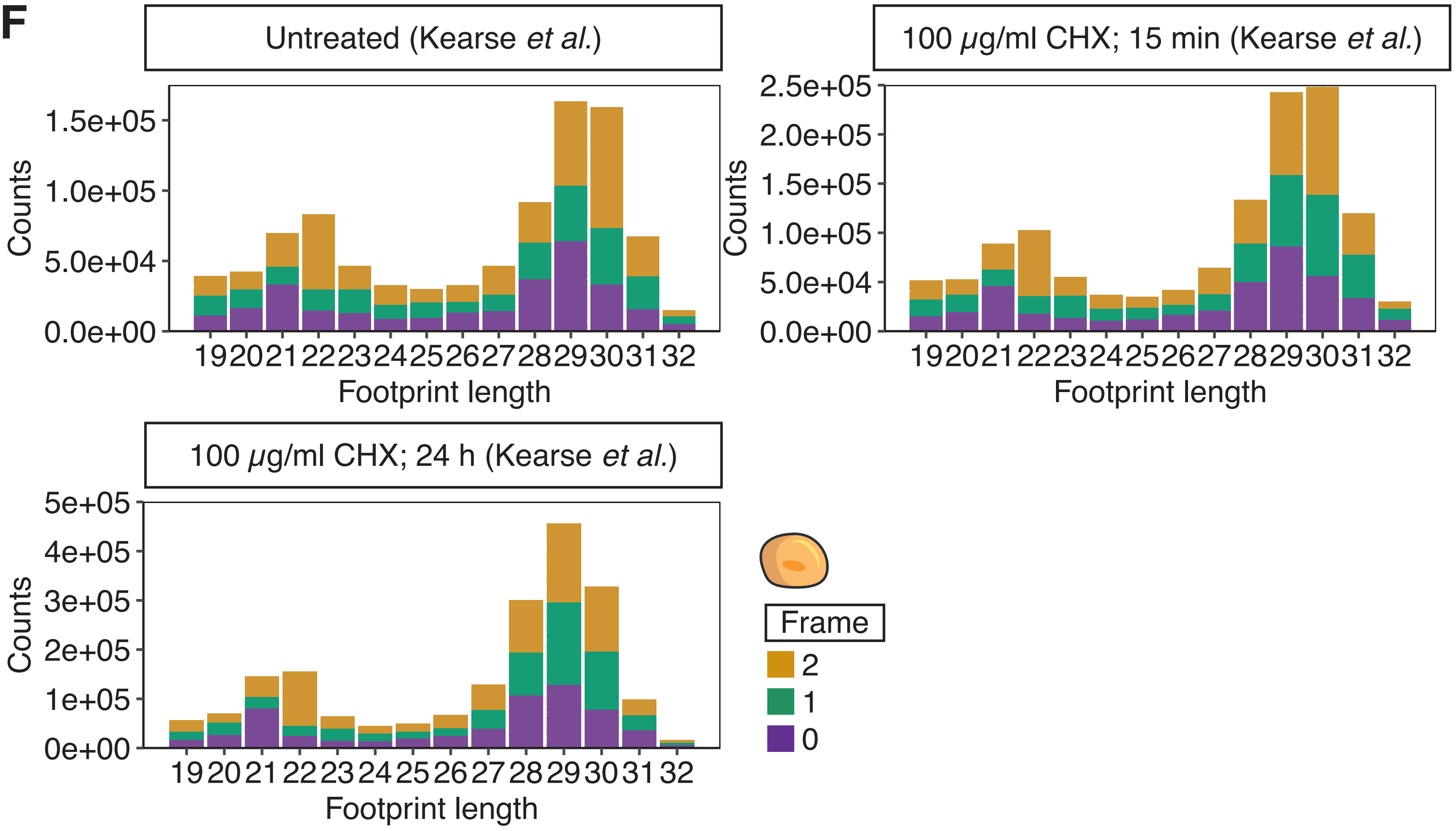


**Supplementary Figure 1**

**Supplementary Figure 1: Nuclease concentration used during lysate digestion determines the dominant footprint length and reading frame.** A) Histograms showing the influence of RNase I concentration on footprint length and reading frame in yeast (+/+) ribosome profiling libraries. Published (1, 2) yeast libraries and the libraries using higher RNase I concentration were generated from long (28-30 nt) footprints using similar protocols. Plots show representative samples. B) Total number and frame of the 5' ends of ribosome footprints of the most common length in a weakly and a strongly digested sample mapped to all yeast coding sequences. 0 matches the first nucleotide of the start codon. The inset shows a magnification to visualize the amount of out of frame reads. C) Same as A) for HEK 293T (+/+) libraries (28-32 nt footprints). D) Percentage of reads mapped to rRNA in HEK 293T libraries prepared using different RNase I concentrations. E) Same as A) for HEK 293T cells treated with different CHX concentrations and incubation period. F) Same as A) for HeLa cells treated with 100 µg/ml CHX for 15 min or 24 h (3). The reading frame is indicative by color: 0 (purple), 1 (green) and 2 (yellow).

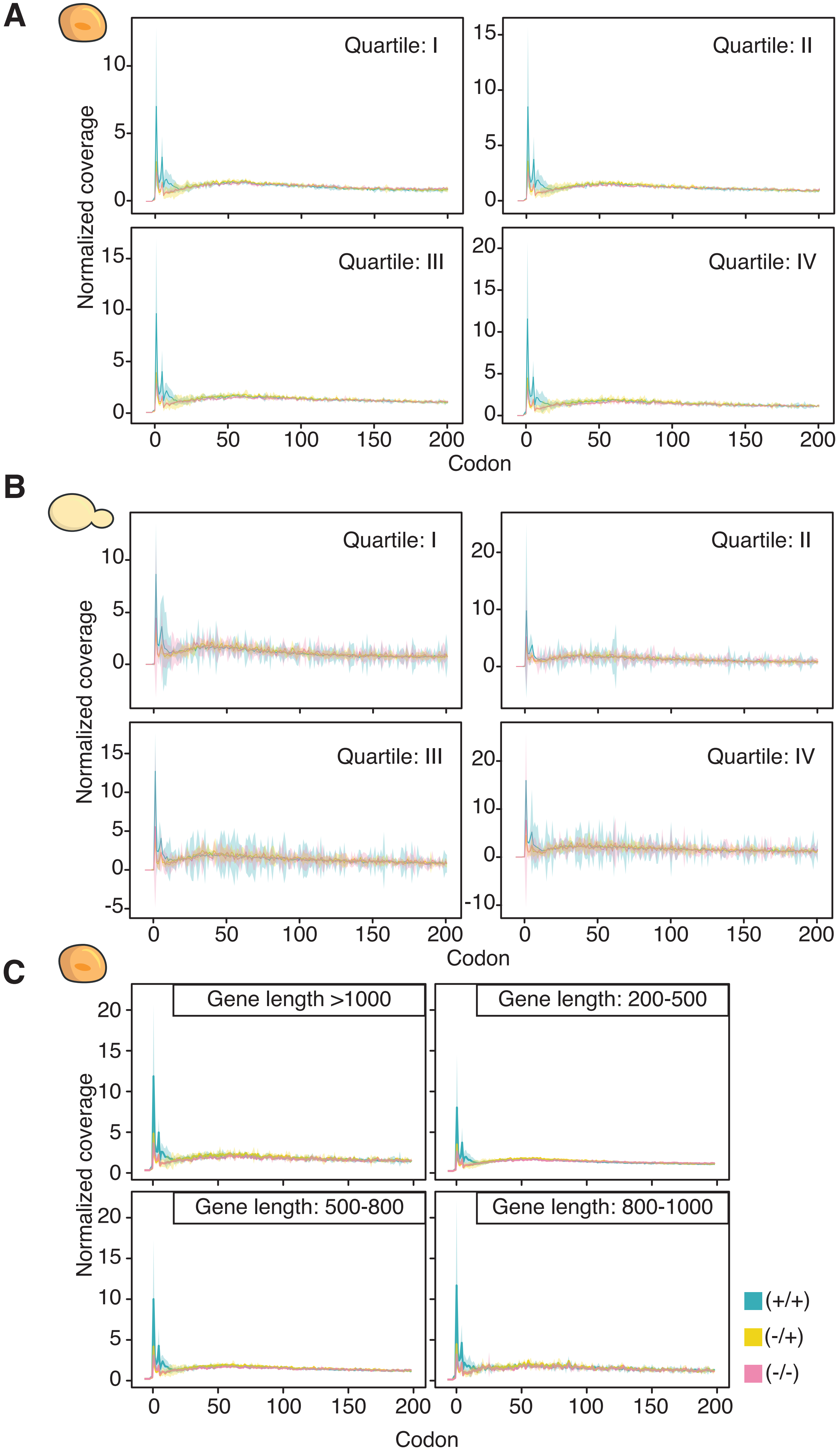


**Supplementary Figure 2**

**Supplementary Figure 2: Ribosome density across ORFs is independent of translation levels and gene length.** A and B) Transcriptome-wide ribosome coverage of human and yeast ORFs divided into quartiles based on translation levels (quartile I: 25% most expressed genes, quartile II: 26%-50% most expressed genes, etc.). C) Transcriptome-wide ribosome coverage of human ORFs divided into four groups based on gene length. Shaded areas represent confidence intervals (n=3). CHX-treatment conditions are indicated by color (+/+) (green), (-/+) (yellow), (-/-) (pink).

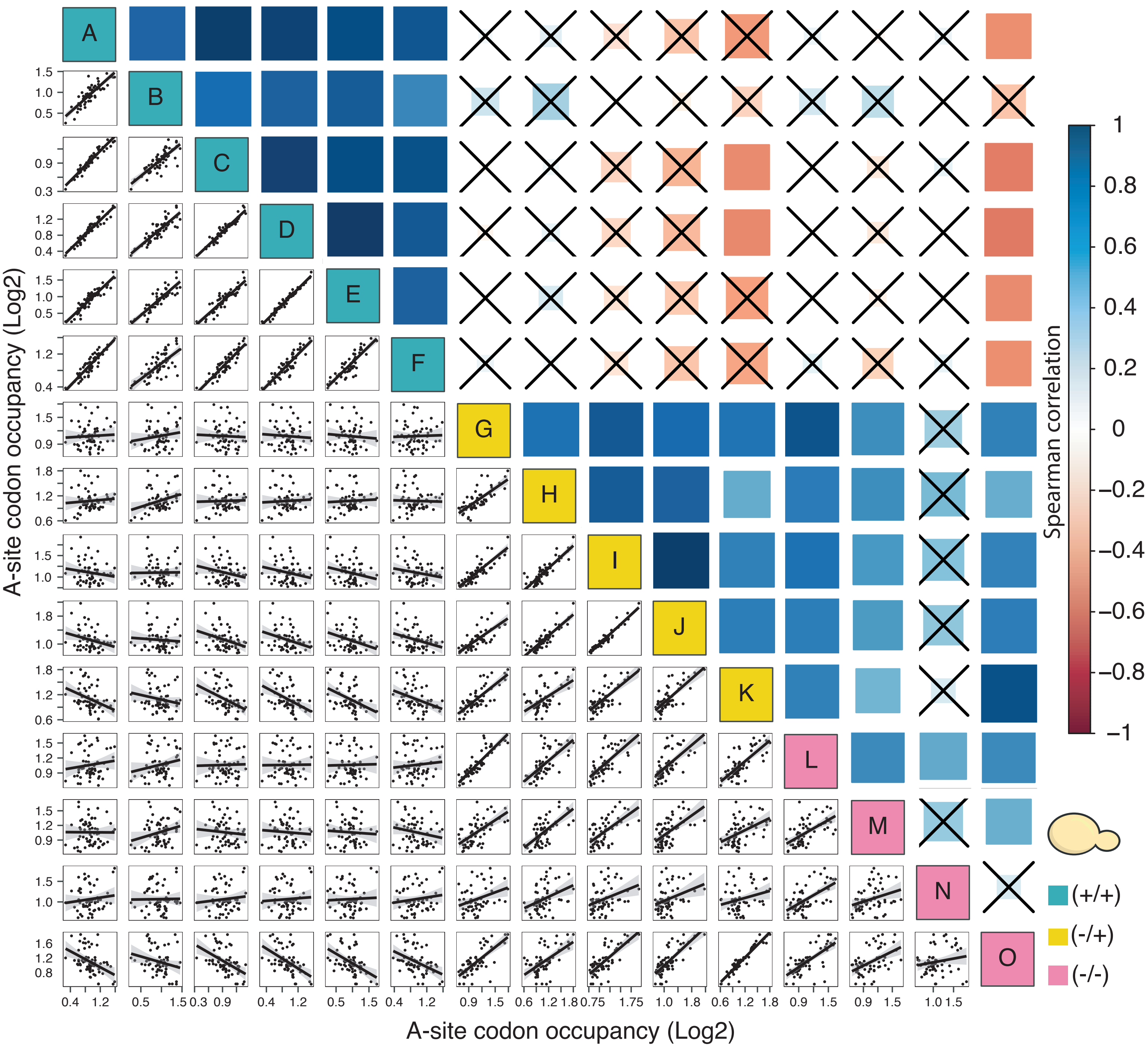


**Supplementary Figure 3**

**Supplementary Figure 3: A-site codon occupancy in yeast is altered by *in vivo* treatment with CHX.** Spearman correlations of A-site ribosome occupancy (compare Figure 3B) across different wild-type ribosome profiling datasets using multiple methods of inhibitor treatment from this study and others (1, 4, 5, 11-16). Size of the box indicates p-value. Correlations with a p-value > 0.05 are crossed out. A: This study; B: (1); C: (4); D: (6); E: (7); F: (5); G: This study; H: (4); I: (5); J: (7); K: (8); L: This study; M: (1); N: (7); O: (8).

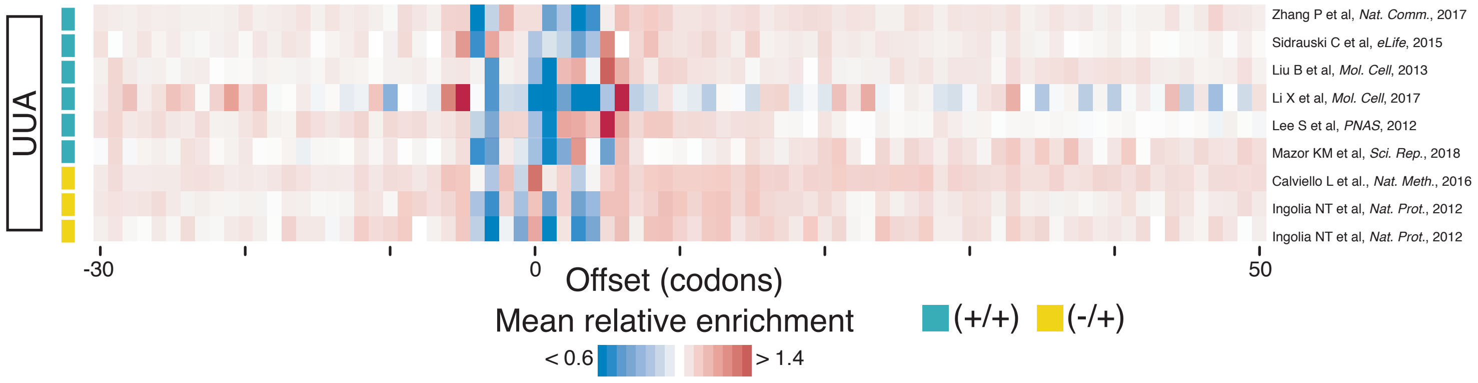


**Supplementary Figure 4**

**Supplementary Figure 4: CHX pre-treatment of vertebrate samples does not alter the ribosome occupancy downstream of rare codons.** Transcriptome-wide ribosome enrichment profiles of published human ribosome profiling datasets (9, 10, 11-16) around the UUA codon with different CHX-treatment methods according to Hussmann and colleagues (33).

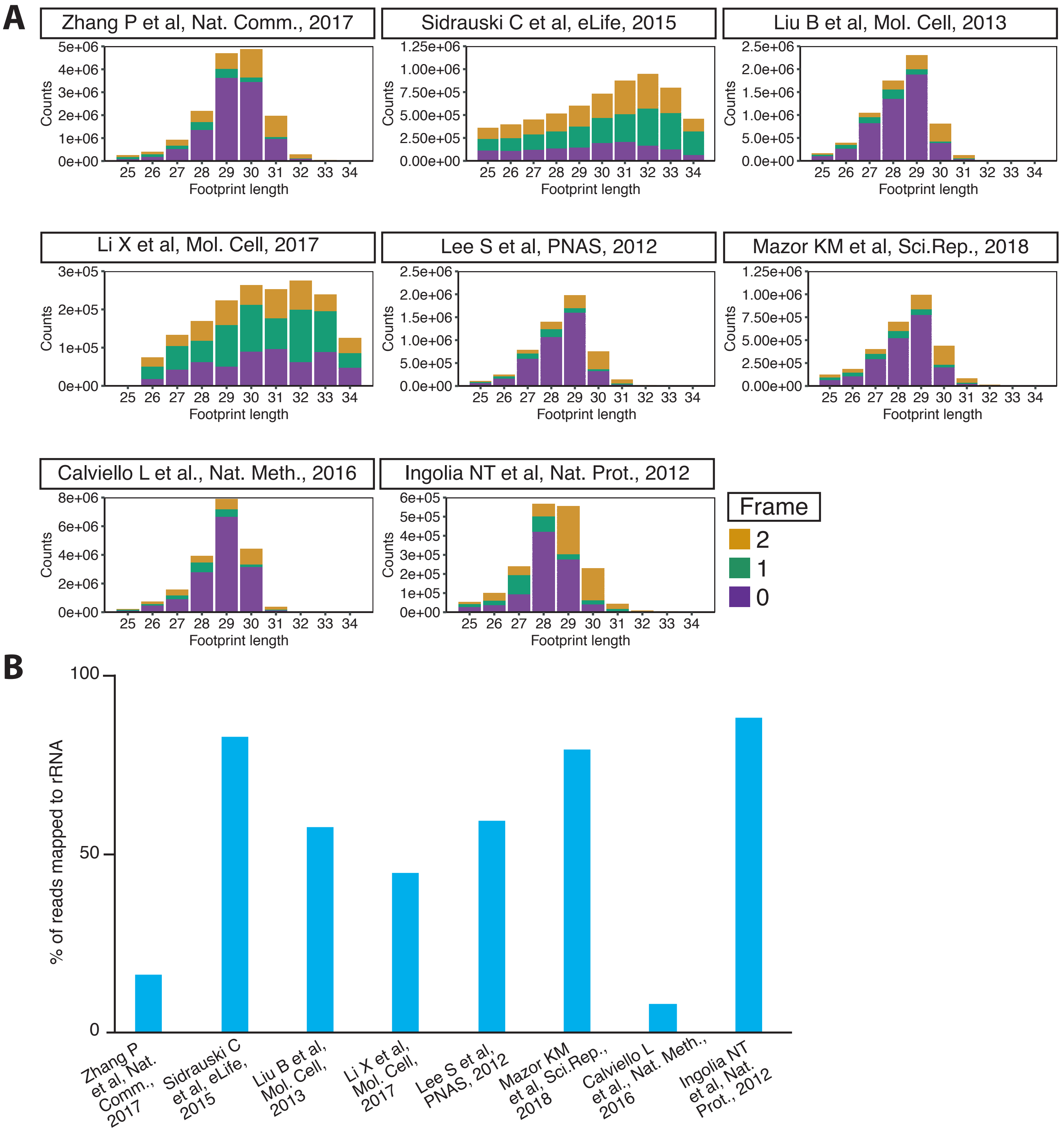


**Supplementary Figure 5**

**Supplementary Figure 5: Differences in footprint length, reading frame and rRNA contamination in published datasets.** A) Histograms showing the footprint length and the reading frame in published human libraries (9, 10, 11-16). The reading frame is indicative by color: 0 (purple), 1 (green) and 2 (yellow). B) Percentage of reads mapped to rRNA in the datasets used in A).
